# A Whole-Brain Computational Modeling Approach to Explain the Alterations in Resting-State Functional Connectivity during Progression of Alzheimer’s Disease

**DOI:** 10.1101/076851

**Authors:** Murat Demirtaş, Carles Falcon, Alan Tucholka, Juan Domingo Gispert, José Luis Molinuevo, Gustavo Deco

**Author notes:** **Corresponding authors**: Gustavo Deco. Computational Neuroscience Group. Universitat Pompeu Fabra.

## Abstract

Understanding the mechanisms behind Alzheimer’s disease (AD) is one of the most challenging problems in neuroscience. Recent efforts provided valuable insights on the genetic, biochemical and neuronal correlates of AD. The advances in structural and functional neuroimaging provided massive evidence for the AD related alterations in brain connectivity. In this study, we investigated the whole-brain resting state functional connectivity (FC) and variability in dynamic functional connectivity (v-FC) of the subjects with preclinical condition (PC), mild cognitive impairment (MCI) and Alzheimer’s disease (AD). The synchronization in the whole-brain was monotonously decreasing during the course of the progression. However, only in the AD group the reduced synchronization produced significant widespread effects in FC. Furthermore, we found elevated variability of FC in PC group, which was reversed in AD group. We proposed a whole-brain computational modeling approach to study the mechanisms behind these alterations. We estimated the effective connectivity (EC) between brain regions in the model to reproduce observed FC of each subject. First, we compared ECs between groups to identify the changes in underlying connectivity structure. We found that the significant EC changes were restricted to temporal lobe. Then, based on healthy control subjects we systematically manipulated the dynamics in the model to investigate its effect on FC. The model showed FC alterations similar to those observed in clinical groups providing a mechanistic explanation to AD progression.

## Introduction

Alzheimer’s disease (AD), being the most prevalent dementia, became a major concern in developed countries as a consequence of increasing life expectancy (Blennow, de Leon, and Zetterberg 2006; Plassman et al. 2007). During the past two decades advancements in genetics, neurobiology and neuroimaging techniques allowed researchers to study the mechanisms behind the underlying causes of AD. In particular, resting state functional Magnetic Resonance Imaging (rs-fMRI) became a useful tool to study the alterations in brain activity of AD patients as well as many other clinical conditions. In the absence of a task, rs-fMRI examines the spontaneous fluctuations of the BOLD signal (Blood-Oxygenation Level Dependent) to identify spatially distributed networks of temporally-synchronized regions. Using seed-based functional connectivity (FC), independent component analysis (ICA) and graph theoretical approaches, recent research on rs-fMRI in AD revealed specific patterns of connectivity associated to the disease (Brier, Thomas, and Ances 2014; Dennis and Thompson 2014; Filippi and Agosta 2011).

The studies that used seed-based approach showed widespread decreases in hippocampal (Allen et al. 2007; W. Li et al. 2012; Wang et al. 2006) and posterior cingulate functional connectivity (Bai et al. 2011; Zhang et al. 2009) in AD. Also, they reported increased FC between prefrontal cortex and hippocampus (Wang et al. 2006), and posterior cingulate (Bai et al. 2011; Zhang et al. 2009) in AD. The increased connectivity regarding prefrontal cortex was interpreted as a compensation mechanism during the initial stages of the disease (Dickerson et al. 2004; Filippi and Agosta 2011; Sanz-Arigita et al. 2010). The studies that used ICA showed decreased activation of default mode network (DMN) (Agosta et al. 2012; Koch et al. 2010; Qi et al. 2010; Sorg et al. 2007) and increased activation of fronto-parietal network (FPN) (Agosta et al. 2012). Various other studies found impaired deactivation of DMN during task in AD and dementia (Celone et al. 2006; Greicius et al. 2004; Lustig et al. 2003; Petrella et al. 2007; Rombouts et al. 2009; Rombouts et al. 2005).

Some studies showed evidence for a prolonged phase of “preclinical AD” in which amyloid-β (Aβ42) plaques accumulate decades before the onset of the first disease symptoms (Price and Morris 1999). The relationship between DMN and AD was further confirmed by the overlap between the spatial pattern of the DMN and that of Aβ42 accumulation that happens in this preclinical phase of AD (Buckner, Andrews-Hanna, and Schacter 2008; Hedden et al. 2009). However, in AD there is a need for biomarkers that link early molecular alterations with later functional manifestations. In this regard, rs-fMRI has been proposed as a promising candidate to bridge this gap (Barkhof, Haller, and Rombouts 2014). In this regard, several studies using rs-fMRI have supplied supporting evidence of a decreased DMN connectivity in cognitively normal individuals with augmented cerebral amyloid load (Sheline, Raichle, et al. 2010; Hedden et al. 2009; Oh et al. 2011). In addition to Aβ42, altered functional connectivity in the DMN have also been associated to abnormal levels of phosphorylated Tau181 (p-tau) in the cerebrospinal fluid (CSF) (Wang et al. 2013) as well as the ratio Aβ42/p-tau, both of which constitute well-established markers of disease progression (X. Li et al. 2013). Indeed, reduced DMN functional connectivity has been reported in amyloid-free carriers of at least one copy of the APOE4 allele, the strongest genetic risk factor for AD (Sheline, Morris, et al. 2010), suggesting that differences in functional connectivity might even precede amyloid deposition (Sheline and Raichle 2013). A recent review emphasized that an integrative approach between connectivity and biomarkers is needed (Ramirez et al. 2014). Furthermore, the structure-function relationship is more complex that it appears (Filippi and Agosta 2011).

In this study, we investigated the grand average FC and the variability in FC (v-FC) across time in Alzheimer’s disease. Grand average FC refers to the conventional measure that quantifies the temporal correlation between brain regions based on Pearson’s correlation coefficient. Variability of FC, on the other hand, takes into account the time dependent variations (standard deviation) of these temporal correlations. For ease of interpretation, to study the alterations during the progression of Alzheimer’s disease we used nodal strengths (i.e. mean connectivity of each individual region) of FC and v-FC. We compared the nodal FC and v-FC strengths of healthy control group (HC) to those of the preclinical (PC), mild cognitive impairment (MCI) and the Alzheimer’s disease patients (AD) groups. Then, we proposed a whole-brain computational model that provided a mechanistic model for the neuronal manifestation of the disease. The model described the BOLD signals of each region with coupled non-linear differential equations that alternates between random fluctuations and sustained oscillations depending on the bifurcation parameter and the interactions between brain regions. As DSI/DTI-based measures cannot capture the biophysical properties (i.e. synaptic conductance, neuronal excitability, time scale…etc.) of the links, we estimated the optimal parameters that aggregate these features (i.e. Effective Connectivity) for each subject. Being specific, effective connectivity (EC) reflects the actual connectivity behind the observed FC. Moreover, we studies the association between FC, v-FC and EC strengths and CSF biomarkers such as Aβ42 and t-tau. Finally, by conducting a computational experiment, we investigated the role of the interplay between random fluctuations and sustained oscillations on the progression of AD.

## Materials and Methods

### Subjects

A total of 109 participants (58 HC, 12 PC, 23 MCI and 16 AD) were recruited at the Alzheimer’s disease and other cognitive disorders unit, from the Hospital Clinic of Barcelona. All subjects underwent clinical and neuropsychological assessment, MRI scanning and were submitted to a lumbar puncture to quantify the content of Aβ42, p-tau and p-tau in CSF. CSF biomarker quantitation was done at the local laboratory by means of ELISA (Enzyme-Linked ImmunoSorbent Assay kits, Innogenetics, Ghent, Belgium). An interdisciplinary clinical committee formed by two neurologists and one neuropsychologist established the diagnoses. HC and PC presented no evidence of cognitive impairment on any of the administered neuropsychological tests, but PC presented an abnormal level of CSF-Aβ42 (below 500pg/ml). MCI and AD presented signs of dementia. MCI patients had an objective memory deficit, defined as an abnormal score on the total recall measure of the Free and Cued Selective Reminding Test (FCRST) (over 1.5 x Standard Deviation), impairment on one or more of the other cognitive tests or preserved activities of daily living, as measured by the Functional Activities Questionnaire (FAQ score <6). The NINCDS-ADRDA criteria were applied for probable AD diagnosis (Jack et al. 2011), taking into account clinical information and objective measures derived from the FAQ and neuropsychological results. AD patients were all in the mild stages of the disease (Global Deterioration Scale = 4). Diagnostic classification was made independent of CSF results. The study was approved by the local ethics committee and all participants gave written informed consent to participate in the study. Genomic DNA was extracted from peripheral blood of probands using the QIAamp DNA blood minikit (Qiagen AG, Basel, Switzerland). Apolipoprotein E genotyping was performed by polymerase chain reaction amplification and HhaI restriction enzyme digestion. Average demographic characteristics of the four diagnostic groups are shown in Table 1.

**Table 1.** Demographics

|  | HC | PC | MCI | AD |
| --- | --- | --- | --- | --- |
| Age | 60.72 (6.99) | 69.00 (7.62) | 69.73 (7.77) | 65.00 (9.98) |
| Sex | 37 (M), 20 (F) | 9 (M), 3 (F) | 14 (M), 9 (F) | 9 (M), 7 (F) |
| APOE carriers | 0.15 | 0.42 | 0.52 | 0.50 |
| Biomarker Index | 0.44 (0.26) | 1.09 (0.55) | 1.50 (0.42) | 1.52 (0.43) |
| AL | 27.23 (6.92) | 21.25 (6.59) | 8.95 (7.14) | 6.23 (4.04) |
| AT | 43.14 (4.44) | 38.42 (7.06) | 21.77 (12.95) | 18.69 (12.48) |
| RDL | 10.43 (2.34) | 8.58 (3.18) | 2.55 (3.20) | 2.00 (2.52) |
| RDT | 14.84 (1.26) | 13.33 (2.46) | 6.32 (5.25) | 5.69 (4.87) |

### Image acquisition and preprocessing

Subjects were examined on a 3T MRI scanner (Magnetom Trio Tim, Siemens, Erlangen, Germany) at the image core facilities of IDIBAPS (Barcelona, Spain). MRI session included a high-resolution three-dimensional structural T1-weighted image (sagittal MPRAGE; TR = 2300ms, TE = 2.98ms; matrix size=256 × 256 x 240; isometric voxel 1×1×1 mm^3^), a ten minute resting state fMRI (rs-fMRI; 300 volumes, TR=2000ms, TE=16ms, 128×128×40 matrix, voxel size=1.72×1.72×3 mm^3^) and two DTI (30 nocollinear directions with a b value=1000 s/mm^2^ and one b0; TR=7700ms, TE=89ms; matrix size=122×122×60; voxel size 2.05×2.05×2 mm^3^).

The pre-processing pipeline of rs-fMRI consisted in the slice-timing correction, the realignment and re-slice, smoothing with a Gaussian kernel (FWHM=5mm), second order de-trending and regressing out Volterra expanded parameters of movement (24 parameters), mean white matter (WM) signal, cerebro-spinal fluid (CSF) mean signal and nulling regressors (Lemieux et al. 2007). The quality criteria to consider a volume wrong and to override it by a nulling regressor, was that its correlation coefficient (cc) with the mean image of its series were beyond three standard deviations (cc < 0.991) from the mean cc of all the images from all subjects to their corresponding mean image (mean cc=0.995). No subjects presented more than 15% of bad volumes, being the average percentage of bad volumes of 1.6% and the standard deviation of 3.7%. To obtain the time series of each region from the Anatomical Automatic Labeling (AAL) atlas (Tzourio-Mazoyer et al 2002), AAL atlas was adapted to every subject native space by co-registering it to the T1 structural image by mean of ANTS (UPENN, UVA and UIowa, USA; http://stnava.github.io/ANTs/). AAL maps in native space were resliced to fMRI resolution using nearest neighbor interpolation and masked with the gray mater (GM) mask. GM mask was constructed for every subject from the tissue probability maps resulted from segmentation of T1 images. The mask was formed by those voxels whose probability of belonging to GM was bigger than the probability of belonging to any other tissue. GM masks were dilated one voxel to include edges and to fill noise-related small gaps and, finally, resliced to fMRI resolution. Time series were obtained by averaging the fMRI signal in the each area of the GM-masked AAL atlas in native space (Tzourio-Mazoyer et al. 2002). The software used for the whole fMRI pre-processing, apart from the above mentioned ANTS, was a homemade MATLAB (Mathworks, Sherborn, MA, USA) script mostly formed by functions from SPM package (Wellcome Trust Center for Neuroimaging; UCL, UK; http://www.fil.ion.ucl.ac.uk/spm/).

Diffusion weighted images (DWI) were first corrected for eddy current distortions using FMRIB Software Library (FSL) package (Jenkinson et al. 2012). We denoised resulting data using the overcomplete local PCA method described in (Manjón et al. 2013). Similarly, T1-weighted image were denoised using a non-local mean filter (Coupe et al. 2008) and then corrected for the usual acquisition bias with the N4 method from the Advanced Normalization Tools (ANT) package (Tustison et al. 2010). Anatomical images were then segmented with the Statistical Parametric Mapping (SPM) VBM8 toolbox (Ashburner and Friston 2000) to create grey matter (GM), white matter (WM) and cerebro-spinal fluid (CSF) probabilistic maps. Bias-corrected T1 images were then co-registered to the non-gradient diffusion image and to the MNI template using respectively ANT’s elastic and symmetric method (Avants et al. 2011). Brain regions of the Anatomical Automatic Labeling (AAL) template (Tzourio-Mazoyer et al. 2002) were then resampled to the anatomical and diffusion space of each subject. Finally, FSL’s Bedpostx and Probtrackx tractography was performed with default parameters on AAL regions, except the cerebellum, resulting in a 90x90 connectivity matrix.

## Whole-Brain Connectivity Measures

We quantified the measures of connectivity based on level of synchronization in the BOLD time series of the four diagnostic groups. After defining a narrowband frequency with 0.03Hz window size, we computed Hilbert transform of the narrowband signal. Hilbert transform converts the narrowband signal as a(t)=A(t)cos(φ(t)), where A(t) is the instantaneous amplitude (or envelope), and A(t), and φ(t) is the instantaneous phase. The first and last 20 seconds (10 TR) of the transformed BOLD signal was then removed. The global coherence and metastability of time-series were computed based on the Kuramoto Order Parameter (KOP)(Kuramoto 1986; Shanahan 2010; Cabral et al. 2012; Hellyer et al. 2014): 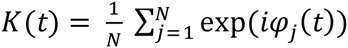, where N is the number of ROIs and φ(t) is the instantaneous phase of each region estimated using Hilbert Transform. The temporal average of Kuramoto Order Parameter defined as the coherence (mean synchronization), while the standard deviation of Kuramoto Order Parameter defined as the metastability (i.e. the variation in synchronization over time). To find the optimal frequency range for band-pass filter, we estimated the group differences in coherence and metastability using 7 distinct frequency onsets varying between 0.01Hz and 0.07Hz (one-way ANOVA). The group differences were optimal at 0.06-0.09Hz narrowband (supplementary figure 1). Functional Connectivity (FC) was computed as the Pearson’s correlations between the time-series of different brain regions. Dynamic Functional Connectivity was computed using a sliding-window analysis approach. We used 2 minutes (60 TR) window size with 20 seconds sliding step size (10 TR). Then, we variability in FC was estimated as the standard deviation of each connection across time. For simplicity we used the mean FC and v-FC of each node characterize the strengths of each node.

### Computational Model

We modeled the whole-brain spontaneous activity using 78 nodes, excluding subcortical regions. Each node was coupled with each other via effective connectivity (EC) matrix. We described the local dynamics of each individual node using normal form of a supercritical Hopf bifurcation. The advantage of this model is that it allows transitions between asynchronous noise activity and oscillations. Where ω is the intrinsic frequency of each node, a is the local bifurcation parameter, η is additive Gaussian noise with standard deviation β, the temporal evolution of the activity, z, in node j is given in complex domain as:

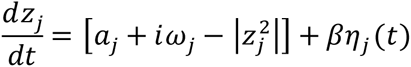

and,

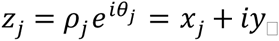

This system shows a supercritical bifurcation at *a_j_=0*. Being specific, if *a_j_* is smaller than 0, the local dynamics has a stable fixed point at *z_j_=0*, and for *a_j_* values larger than 0, there exists a stable limit cycle oscillation with a frequency *f=ω/2π*. Finally, the whole-brain dynamics is described by the following coupled equations:

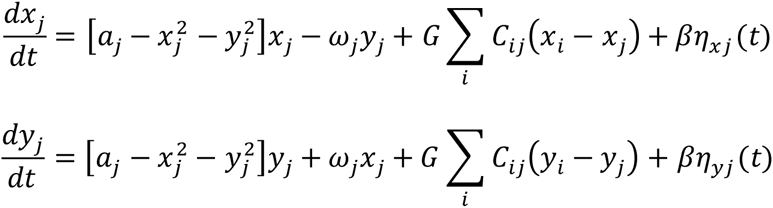

Where *C_ij_* is the Effective Connectivity (EC) between nodes i and j, *G* is the global coupling factor, and the standard deviation of Gaussian noise, *β=0.02*. The simulated activity corresponds to the BOLD signal of each node. The simulations were run for 30000s, sampled at 2s, if not stated otherwise. Both the empirical and simulated BOLD signals were band-pass filtered in narrowband 0.06–0.09Hz, since the group differences in coherence and metastability were optimal in this narrowband. The intrinsic frequency of each node was estimated as the peak frequency in the associated narrowband of the empirical BOLD signals of each brain region.

### Optimization of Effective Connectivity

We implemented a heuristic approach to infer the most likely connectivity matrix (i.e. Effective Connectivity) that explains the empirical functional connectivity. As an initial guess, we started with the anatomical connectivity matrices. First, we adjusted the global coupling parameter (G) to prevent overflow during the optimization procedure, and to ensure the stability of the system of equations. Where *K^sim^* and *K*^*em p*^ are simulated and empirical coherences (average Kuramoto order parameter), we updated global coupling parameter as: 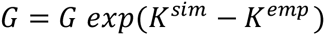, until the desired condition, 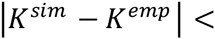 0.1, was satisfied.

We evaluated both zero-time-lag Functional Connectivity (FC^0^) and time-lagged FC (FC^τ^) for both empirical and simulated BOLD signals. Time-lagged FC measure was chosen for two reasons. First, it provides an additional constraint ensuring that the optimal solution is unique. Second, time-lagged correlations allow inference on the directionality of the connections. The defined the distance metric as Euclidean Distance between simulated and empirical FC values for both FC^0^ and FC^τ^:

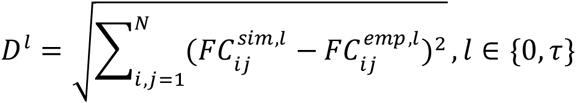

Then, where E is the average error between empirical and simulated FC measures;

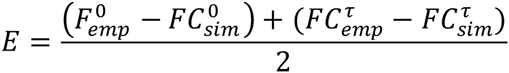

Where *S_ij_* is the anatomical connectivity matrix, N is the number of regions, and Λ denoted inter-hemispheric links; we updated the effective connectivity between i and j=(1,…, N) according to:

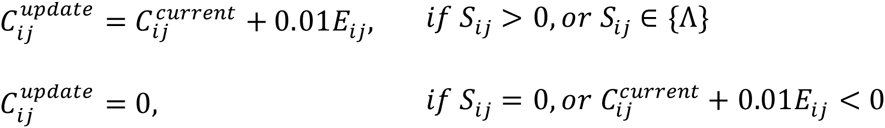

In other words, we updated the EC based on the average error between empirical and simulated FC measures for non-zero connections and inter-hemispheric connections. Negative weights were not allowed, and they all set to zero during the update procedure. We accepted the total distance between empirical and simulated FC measures, D^T^=D^0^+D^τ^, for updated EC is lower than the minimum total distance observed during the procedure. We repeated this procedure using 100 iterations, and the best solution (minimum D) was considered as EC for a given subject. The entire procedure was also repeated for bifurcation parameter *a={-0.1, -0.05, 0, 0.05, 1}*. The best fit was achieved at a=-0.05 (supplementary figure 2A). Finally, the joint strengths of EC were computed to quantify the overall strength of each node.

Given the inferred EC matrices, we disrupted the dynamics in healthy controls based on the bifurcation parameter, a, of the supercritical Hopf Normal Model. First, we computed the FC of the healthy controls at a=-0.05, where the optimal similarity between simulated and empirical values was observed. Then, we decreased the bifurcation parameter by 0.001 at each iteration, spanning values between -0.05 and -0.15. The nodal strengths of the FC and the variability of FC were computed for each value and quantified the alterations as FC^a^ – FC^-0.05^.

The predicted alterations in healthy controls were then compared to the empirical alterations observed in clinical populations as FC^clinical^ – FC^HC^ (clinical={PC,MCI,AD}. The similarity was quantified as Euclidean distance between predicted and empirical alterations.

### Statistical Analyses

The group comparisons (for coherence, metastability, FC strength, v-FC strength, EC strength) were done using permutation t-test (1000 permutations), and multiple comparisons were corrected using FDR approach with Benjamini&Hochberg algorithm if necessary (Hochberg and Benjamini 1990). Prior to group comparisons, we regressed out subject’s age from each measure. The networks were visualized using BrainNet Viewer toolbox in Matlab (Xia, Wang, and He 2013). Correlations between CSF biomarkers (APOE-4, Aβ-42, and t-tau) and the measures (coherence, metastability, FC strength, v-FC strength, EC strength) were estimated as the partial correlations controlled for age, gender and education level. APOE was quantified as the carriers and non-carriers of the gene.

## Results

### Alterations in Global Connectivity Measures

We compared the mean (coherence) and standard deviation (metastability) of the Kuramoto order parameter (synchronization) of PC, MCI, and AD groups with the healthy controls (HC) using permutation t-test (10000 permutations, p-value < 0.05). The results showed a monotonous decrease in both measures along with the progression of the disease (Figure 2). The coherence and metastability of the AD group were significantly different than HCs (coherence: T-statistic=3.4, p-value < 0.01; metastability: T-statistic=3.27, p-value < 0.01) and PCs (coherence: T-statistic=2.77, p-value < 0.05; metastability: T-statistic=2.84, p-value < 0.05). We found no significant differences in the rest of the clinical populations.

**Figure 1.**
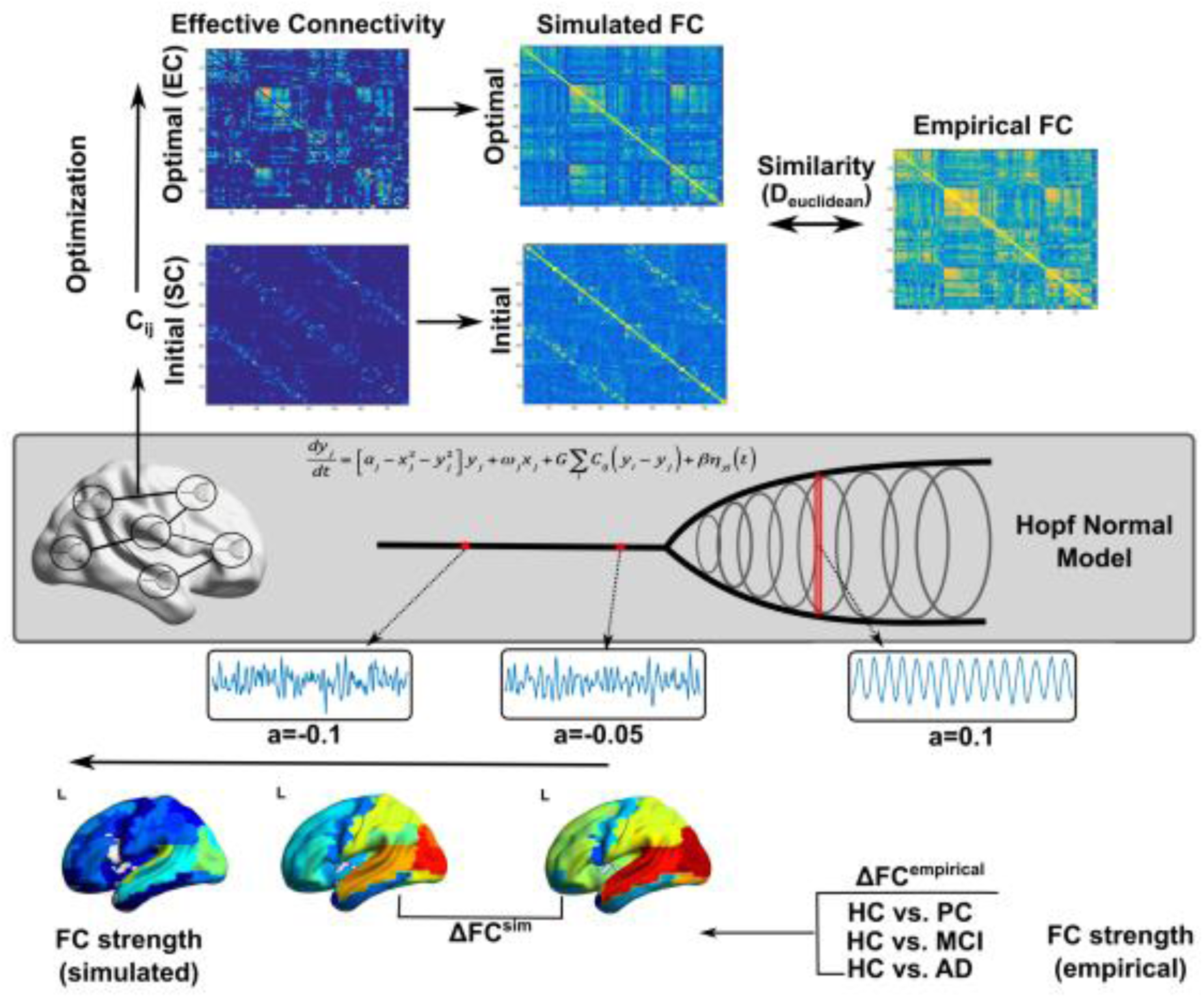
Overview of the model. Top panel illustrates the optimization procedure. Middle panel illustrates Hopf normal model. Bottom panel illustrates the computational experiment.

**Figure 2.**
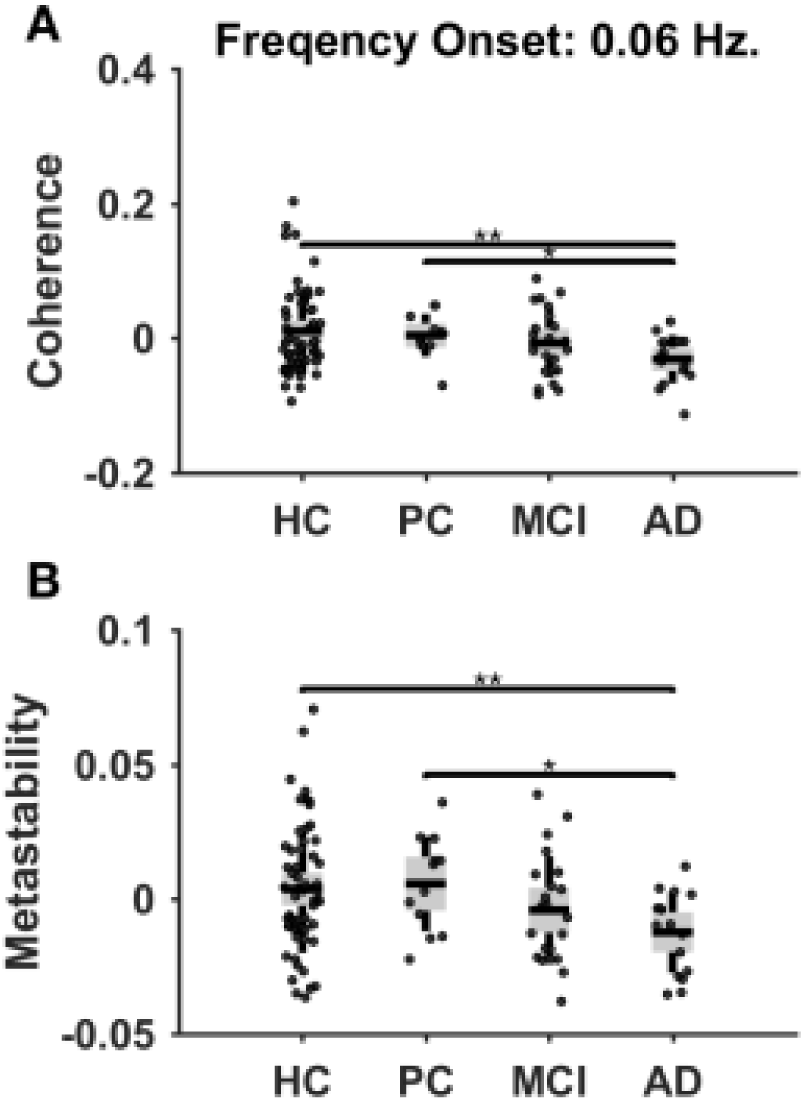
Group differences in coherence (average Kuramoto order parameter) and metastability (standard deviation of Kuramoto order parameter) in 0.06-0.09Hz narrowband signal. The comparisons were done using permutation t-test (1000 permuations; * p<0.05, ** p<0.01).

### Alterations in Regional Connectivity Measures

We compared the nodal strengths of functional connectivity (FC), variability of functional connectivity (v-FC), and effective connectivity (EC) of PC, MCI and AD groups with the HCs. The nodal strengths of FC were decreased in the entire brain in PC, MCI and AD groups (Figure 3). AD group exhibited significantly decreased FC strengths widespread across the brain including the hubs in prefrontal and parietal regions (Table 2). The differences were more prominent in right hemisphere. No significant differences were found in FC strengths of PC and MCI groups.

**Figure 3.**
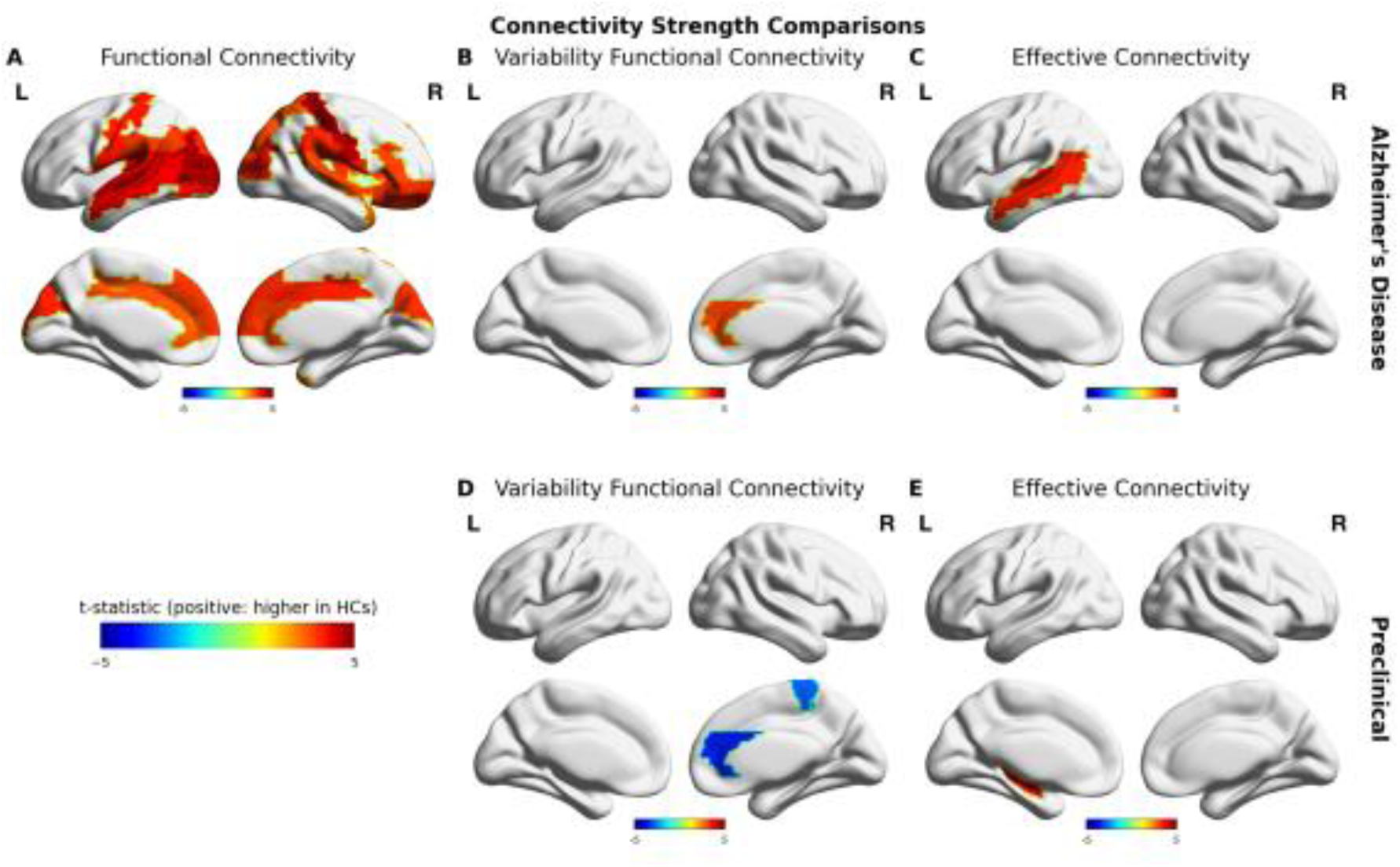
Group differences in nodal strengths of FC, vFC, and EC (columns). Top row shows the comparisons between AD and HC groups, bottom row show comparisons between PC and HC groups. The comparisons were done using permutation t-test (1000 permuations). Colorbars indicate T-statistic. Only significant differences after FDR correction were reported.

**Table 2.** Group comparisons FC strength (T-statistic) FC Strength (AD vs. HC)

| FC Strength (AD vs. HC) |  |  |
| --- | --- | --- |
| ROI | Left | Right |
| PreCG | 1.9468 | 2.8517* |
| SFGdor | 1.8606 | 2.3775* |
| ORBsup | -0.1604 | 3.2579** |
| MFG | 1.0807 | 2.6271* |
| ORBmid | -0.4677 | 3.3205** |
| IFGoperc | 1.3351 | 2.4945* |
| IFGtriang | 2.4744* | 3.0819** |
| ORBinf | 0.6945 | 3.3300* |
| ROL | 3.4757** | 3.4013** |
| SMA | 2.5766* | 1.8211 |
| OLF | -1.1313 | 0.2452 |
| SFGmed | 3.1908* | 3.3281** |
| ORBsupmed | 2.4408* | 2.8573* |
| REC | 0.8826 | 0.4747 |
| INS | 1.7096 | 3.2364** |
| ACG | 2.8703* | 3.1387** |
| DCG | 2.9755** | 3.4686** |
| PCG | 0.2593 | -0.5377 |
| HIP | -0.3394 | 2.1593* |
| CAL | 1.6324 | 1.9620 |
| CUN | 3.4414** | 3.2861** |
| LING | 1.8809 | 2.0415 |
| SOG | 3.3086** | 3.0077* |
| MOG | 4.6237** | 3.5258** |
| IOG | 3.6295** | 2.6303* |
| FFG | 2.9761* | 1.0922 |
| PoCG | 3.5312** | 4.5427** |
| SPG | 1.8369 | 2.8759* |
| IPL | 2.3735* | 1.3246 |
| SMG | 3.2118** | 3.3754** |
| ANG | 3.1634** | -0.0517 |
| PCUN | 2.3624* | 2.5367* |
| PCL | 1.7343 | 2.1213 |
| HES | 3.7915** | 1.8432 |
| STG | 4.0204** | 3.0325** |
| TPOsup | 1.1001 | 1.7304 |
| MTG | 3.9036** | 2.5762* |
| TPOmid | 0.4231 | 2.8015* |
| ITG | 1.2090 | 1.0448 |

\* p < 0.05
\*\* p < 0.01

FC variability strength in right anterior cingulate cortex was increased in PC group (T-statistic=-3.4667, FDR-corrected p-value < 0.05), while it was decreased (T-statistic=3.0661, FDR-corrected p-value < 0.05) in AD group. Right paracentral lobule was also decreased in FC variability strength in PC group (T-statistic=-2.8062, FDR-corrected p-value < 0.05). No significant differences were found in MCI group.

In order to reveal the underlying connectivity alterations, we used whole-brain computational model to infer EC. The average correlation coefficient between empirical and simulated FC was r = 0.81 (std=0.03). The correlation coefficients between empirical and simulated coherence and metastability were r = 0.93 (p-value < 0.001) and r = 0.72 (p-value < 0.001), respectively (supplementary figure 2). We found decreased nodal EC strengths in left hippocampus in PC group (T-statistic=3.5721, FDR-corrected p-value < 0.05) and in left medial temporal gyrus in AD group (T-statistic=3.4651, FDR-corrected p-value < 0.05). No significant differences were found in MCI group.

### Relation to APOE4 and CSF Biomarkers

Coherence showed a significant correlation with Aβ-42 (rho = 0.25, p-value < 0.05), but not t-tau (rho = -0.13, p-value=0.19) for all subjects. However, within clinical groups (i.e. PC, MCI, and AD) coherence was not correlated with Aβ-42 (rho = 0.09, p-value=0.53) and t-tau (rho = -0.01, p=0.96) CSF biomarkers. In contrast, we found no significant correlations between metastability and Aβ-42 CSF biomarker (clinical groups: rho = 0.03, p-value=0.86; all subjects: rho = 0.15, p-value=0.13), but it was significantly correlated with t-tau CSF biomarker (clinical groups: rho = -0.31, p-value < 0.04; all subjects: rho = -0.27, p-value < 0.05).

We also investigated how APOE-4 and CSF biomarkers are associated to FC, vFC and EC strengths (figure 4). CSF biomarker for Aβ-42 exhibited significant correlations with FC strengths of bilateral rolandic operculum (left: rho = 0.25; right: rho = 0.27, p-value<0.05), middle cingulate gyrus (left: rho = 0.23; right: rho = 0.23, p-value<0.05), cuneus (left: rho = 0.26; right: rho = 0.24, p-value<0.05), superior (left: rho = 0.26; right: rho = 0.20, p-value<0.05) and middle (left: rho = 0.33; right: rho = 0.20, p-value<0.05) occipital lobes, and postcentral gyrus (left: rho = 0.27; right: rho = 0.26, p-value<0.05). In left hemisphere Aβ-42 biomarker was significantly correlated with FC strenghts of inferior parietal lobe (rho = 0.24, p-value<0.05), supramarginal gyrus (rho = 0.25, p-value<0.05), angular gyrus (rho = 0.23, p-value<0.05), heschl gyrus (rho = 0.25, p-value<0.05), superior (rho = 0.30, p-value<0.05) and middle (rho = 0.34, p-value<0.05) temporal lobes, in right hemishere significant correlations were observed in precentral (rho = 0.20, p-value<0.05) and paracentral (rho = 0.28, p-value<0.05) lobules, superior (rho = 0.20, p-value<0.05) and middle (rho = 0.23, p-value<0.05) frontal lobes, pars opercularis (rho = 0.24, p-value<0.05), pars triangularis (rho = 0.28, p-value<0.05), and pars orbitalis (rho = 0.22, p-value<0.05) of inferior frontal gyrus. CSF biomarker for t-tau was significantly correlated with FC strengths of left superior (rho = -0.23, p-value<0.05) and inferior (rho = -0.23, p-value<0.05) occipital lobes, left superior parietal lobe (rho = -0.22, p-value<0.05), left paracentral lobule (rho = -0.20, p-value<0.05), right superior (rho = -0.20, p-value<0.05) and middle (rho = -0.24, p-value<0.05) frontal lobes. No significant correlations were observed between FC strength APOE-4 allelle carrier status. Variability of FC strength showed significant correlations with Aβ-42 CSF biomarker in right pars triangularis of inferior frontal gyrus (rho = -0.21, p-value<0.05), right middle cingulate gyrus (rho = -0.26, p-value<0.05), right paracentral lobule (rho = -0.21, p-value<0.05), and right temporal pole (rho = -0.24, p-value<0.05). T-tau CSF biomarker was significantly correlated with FC variability strengths of bilateral inferior parietal lobe (left: rho = -0.20; right: rho = -0.26, p-value<0.05), left superior occipital lobe (rho = -0.21, p-value<0.05), right supramarginal gyrus (rho = -0.24, p-value<0.05), and right paracentral lobule (rho = -0.20, p-value<0.05). APOE-4 allelle carrier status was significantly correlated with FC variability strengths in left superior parietal lobe (rho = -0.22, p-value<0.05), right middle frontal gyrus (rho = -0.20, p-value<0.05), right inferior parietal lobe (rho = -0.23, p-value<0.05), right supramarginal gyrus (rho = -0.23, p-value<0.05) and right temporal pole (rho = 0.30, p-value<0.05).

**Figure 4.**
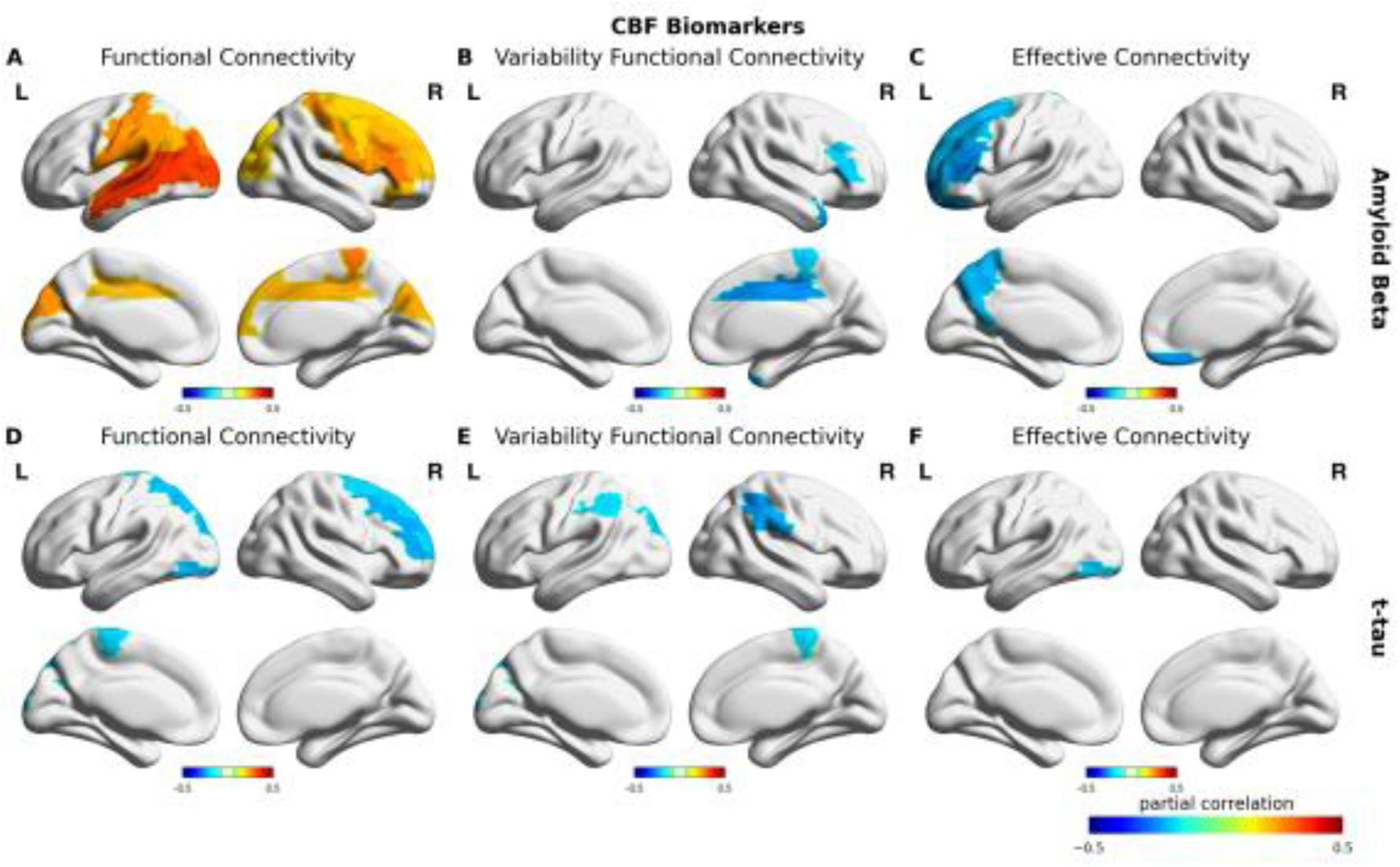
Partial correlations between FC, v-FC and EC (columns) and Aβ-42 (top row) and t-tau (bottom row) controlled for age, gender and education level. Colorbars indicate correlation coefficient (rho). Only significant correlations were reported.

The correlation between EC strength and Aβ-42 CSF biomarker was significant in left superior (rho = -0.21, p-value<0.05) and medial (rho = -0.30, p-value<0.05) frontal gyrus, left superior (rho = -0.22, p-value<0.05) and medial (rho = -0.20, p-value<0.05) orbital frontal gyrus, left pars triangularis of inferior frontal gyrus (rho = -0.25, p-value<0.05), left precuneus (rho = -0.22, p-value<0.05), and right rectus (rho = -0.25, p-value<0.05). EC strength was significantly correlated with t-tau only in left inferior occipital lobe (rho = -0.22, p-value<0.05). No significant correlations were observed between EC strength and APOE-4 allelle carrier status.

### Computational Experiment

Statistical comparisons between groups regarding global (coherence and metastability) and nodal (FC and vFC strengths) measures showed consistent yet complex progression of the alterations in whole-brain connectivity. However, the alterations in EC strength did not show widespread alterations as it was observed in FC strength. For this reason, we conducted a computational experiment to check whether there is a global factor that might explain the progression of the disease. Using the proposed model, we systematically altered the local bifurcation parameter (*a*) in healthy control subjects to study the progression of the disease. Being specific, we disrupted the optimal operating point of the model in favor of noisy dynamics in healthy control subjects and check the effects of this manipulation on simulated FC strengths. The optimal similarity (Euclidean distance) between simulated and empirical FC strength changes were observed at a=-0.065 for PC, a = -0.068 for MCI, and a = -0.084 for AD groups (Figure 5). We found that also the changes in the ratio between mean FC and v-FC in the model were in line with the empirically observed ratios at these optimal local bifurcation parameter values. Furthermore, the correlation coefficients between the simulated and empirical alterations in FC strengths reached up to r = 0.46 (a=-0.088) for PC, r = 0.62 (a = -0.137) for MCI, and r = 0.72 (a = -0.139) for AD (supplementary figure 4-5).

**Figure 5.**
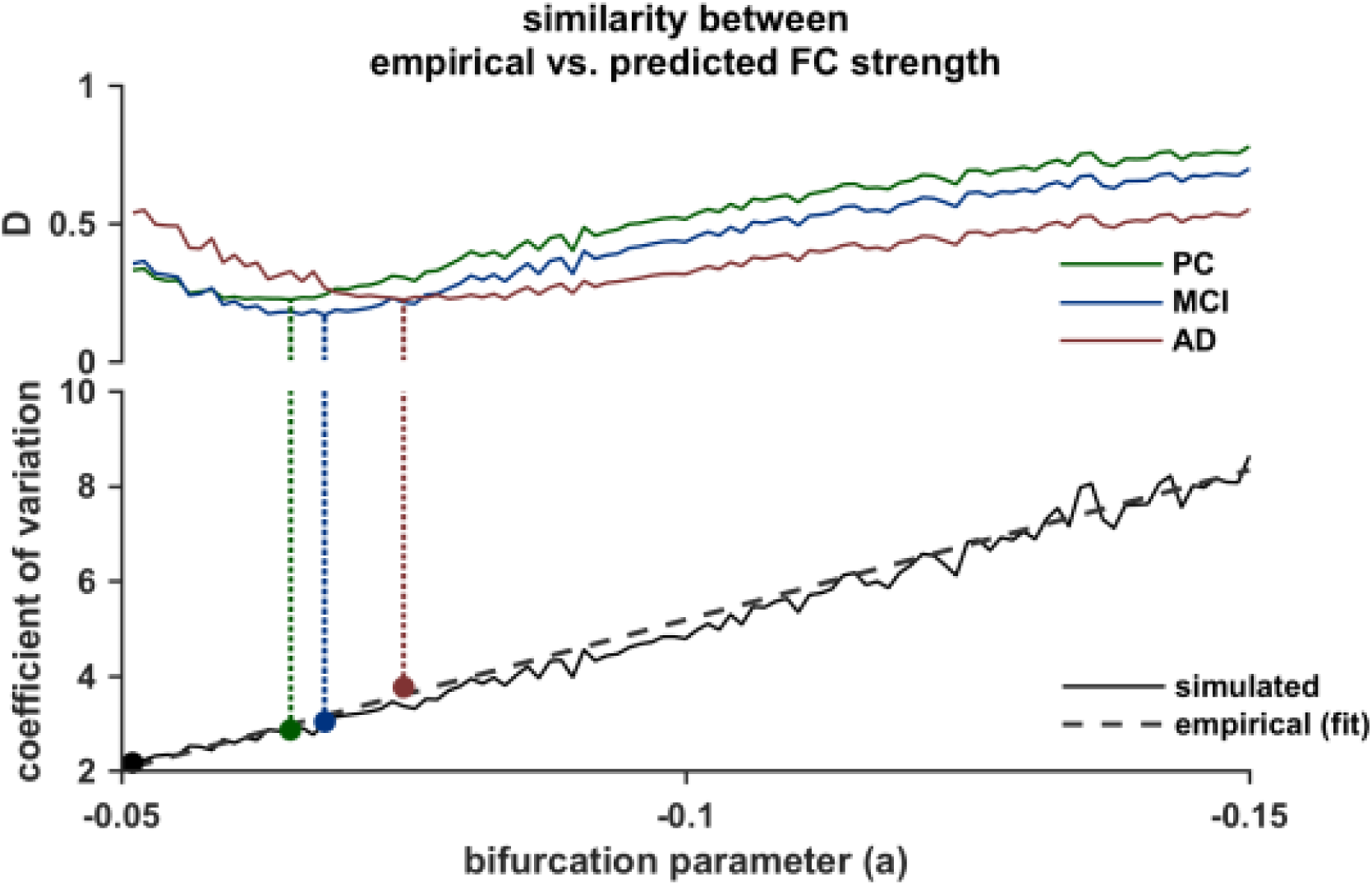
Computational experiment based on altered local bifurcation parameter of healthy control subjects within the range -0.05 and -0.15. The optimal fit for HCs is at a=-0.05. Top: The similarity (Euclidean distance) between FC nodal strength changes in the simulated data (ΔFC^a^- ΔFC^a=-0.05^) and in the empirical data (green: ΔFC^PC^- ΔFC^HC^, blue: ΔFC^MCI^- ΔFC^HC^, red: ΔFC^AD^- ΔFC^HC^). Dashed lines indicate the minimum distance. Bottom: The ratio between grand average v-FC and FC. Circles indicate the empirical values for HCs (black), PC (green), MCI (blue) and AD (red) projected to the optimal fit to the simulated data. Dashed line is the linear fit to the empirical data.

## Discussion

In this paper, we studied the whole-brain functional connectivity in Alzheimer’s disease. We treated the condition as a continuum based on severity comprising PC, MCI and AD. We found that the coherence (average synchronization) and metastability (standard deviation of synchronization) monotonously decrease through this continuum. The results showed gradual decreases in FC as the disease progresses. However, the decrease in FC was only significant in AD group with respect to the HC group.

Furthermore, the higher order statistics of the FC (i.e. variability of FC) provided critical information that might help the characterization of the disease. Increased v-FC in PC group suggests that presence of amyloid plaques without signs of dementia might be linked to abnormal FC dynamics. Therefore, a possible mechanism for the onset of neurodegenerative diseases might be the divergence from the optimal dynamical regime, in which the connectivity still remains intact. We found that the v-FC in anterior cingulate cortex (ACC) was significantly increased in PC group, but decreased in AD group. We speculate that whatever causes elevated variability in ACC in PC group might also be responsible for reduced variability observed in AD group. Ineffective communication in the entire cortex would primarily affect the connectivity hubs in the brain. Therefore, a small perturbation in whole-brain dynamics may spread out and cause the elevated variability in FC. As the perturbation advances to a point where the whole-brain communication collapses, its effect might qualitatively change. Therefore, the dissociation between anterior and posterior parts of DMN, and the hyper-synchronization of several networks that was reported in previous studies might be explained as a network effect in the whole-brain.

Nevertheless, the data driven methods based on FC and v-FC reflect the cumulative effects of complex interactions between brain regions. Therefore, understanding the underlying mechanisms behind FC is non-trivial without model-based approaches. We proposed a computational model to characterize the interplay between coordinated noise and oscillations in the whole-brain. We estimated the optimal EC from the empirical observations to make inference on the causal origins of the FC alterations. Despite the widespread changes in FC in AD, EC revealed that right temporal lobe was the only region where the connectivity was actually altered. Moreover, in PC group EC was decreased in the hippocampus suggesting the role of this region in even there are no signs of memory deficits.

It is important to note that the model-based inference of EC presented in this study should be considered in more abstract terms that are dependent to the computational model. Being specific, the EC reflects a high-level parameter space that lumps various biophysical features of the neural network as well as hemodynamic properties that cannot be accounted by the proposed model. However, the major advantage of this approach is that it allows constructing model-based hypotheses based on critical model parameters, given the EC. In this study, we focused on the influence of the bifurcation parameter on the FC alterations. We showed that the divergence from the bifurcation point (i.e. nodes exhibit noise dynamics) might cause similar alterations observed in the empirical data. This supports the view that perturbations in whole-brain dynamics cause structured alterations in FC. Moreover, the model illustrated slightly increased overall v-FC as observed in PC group, which later decreased as the system diverged more from the bifurcation point. The best correspondence between FC alterations in the simulations and those in the AD group was observed at the point where the v-FC started to drop (supplementary figure 3).

In this study we also addressed the influence of the genetic and the CSF biomarkers. We found that although coherence was correlated with Aβ-42 and t-tau, these correlations were driven by the contrast between healthy controls and clinical populations, as the correlations disappeared among clinical populations. On the other hand, t-tau CSF biomarker was significantly correlated with metastability both for all subjects and clinical groups. Therefore, metastability might be a robust measure that reflects the cognitive function of the brain as it was suggested before (Deco and Kringelbach 2014). The correlations between CSF biomarkers and FC were consistent with the alterations observed in AD group. Furthermore, these results suggest that Aβ-42 is more relevant to the presence of connectivity alterations, but not the progression of the disease. This is consistent with the related studies stating that amyloid-β associated abnormal DMN connectivity in elders without dementia (Sheline, Raichle, et al. 2010; Sperling et al. 2009). Furthermore, the results support the findings that although amyloid-β has a major impact on the neurodegeneration, it is functionally less relevant than tau tangles (Yoshiyama, Lee, and Trojanowski 2013). Another study also discussed the role of Aβ-42 as a necessary but not sufficient factor for AD related dementia (Musiek and Holtzman 2015). Our results are in line with this conclusion.

Nevertheless, the model we propose to infer EC have some limitations. The most critical disadvantage rises from the optimization procedure. Given a highly non-linear model, the use of well-established techniques such as Bayesian inference becomes computationally infeasible. Heuristic approaches to infer EC are difficult to validate given the complexity of the model and the available data. We addressed some of these problems by constraining the parameter space to non-zero connections in structural connectivity (SC) and limiting the possible values between 0 and 1.

Our study showed that the neuronal basis of the transition between neurodegeneration and cognitive impairment might be understood by investigating the abnormalities in FC variability. We speculate that at this stage the outcome of the neuronal atrophy might be determined by peripheral mechanisms (e.g. involvement of neurotransmitters, nonneuronal cells…etc.) that modulate the temporal aspects of the connectivity. Further studies might focus on the local circuitry and biophysical mechanisms behind the FC variability. Nevertheless, it is often non-trivial task to propose biophysically realistic models to study brain function in mesoscopic scale. We believe that as EC provides a quantitative measure to describe the causal origins of whole-brain connectivity, it can provide useful insights to propose better models.

